# Transcriptomic divergence of network hubs in the prenatal human brain

**DOI:** 10.1101/2025.05.26.656237

**Authors:** S. Oldham, G. Ball

## Abstract

Connections in the human brain are not uniformly distributed; instead, a dense network of long-range projections converges on highly connected hub regions located in transmodal, paralimbic, and association cortices. Hub connectivity is strongly influenced by genetic factors but the molecular cues guiding the foundation of these structures remain poorly understood. Here, we combined high-resolution diffusion MRI tractography from 208 term-born neonates with spatially resolved prenatal gene expression data to investigate the molecular correlates of network hub formation at mid-gestation. We identified robust hub architecture in the neonatal connectome and mapped these structural hubs to corresponding cortical regions in the μBrain prenatal digital brain atlas. Transcriptomic analysis revealed differential gene expression in network hubs prior to the time of birth, with genes positively associated with hub status supporting the establishment of early neuronal circuitry and predominantly expressed in the transient subplate and intermediate zones. Hub genes were expressed by excitatory neurons, including subplate neurons and intratelencephalic projection neurons in deep cortical layers, and overlapped with markers of cortical expansion and interhemispheric connectivity in adulthood. Our study identifies prenatal transcriptomic signatures of network hubs in the neonatal human brain, demonstrating how early gene expression programs can shape brain network connectivity from mid-gestation.

## Introduction

Higher order brain functions are founded upon distributed neural activity over anatomical networks.^1–4^ Studies of brain network organisation have revealed a set of fundamental properties that span neuronal networks across species and scales.^1,5–12^ These include: a hierarchical modular configuration, a central core of densely connected, high-degree ‘hub’ regions and, in terms of connection length, near minimal wiring costs.^5,10,13–16^ In the human cortex, hub nodes with extensive, distributed connectivity are located in transmodal, paralimbic and association areas.^17–20^ The central positioning of hubs within brain networks supports integration across functional sub-systems; a framework that is supported by experimental tracer studies in other mammalian species.^4,16,21–24^

Connections between brain regions are unevenly distributed.^25–27^ A disproportionate number of connections form between hubs to forge a central nexus, or ‘rich club’.^5,19,28–32^ Compared to non-hub regions, cortical network hubs have higher metabolic energy demands and increased blood flow.^33,34^ The high metabolic demand, coupled with dense interconnectivity, of core hubs points to a significant energetic cost of rich club ordering in brain networks.^5,10,30,35^ The cost of a rich network core is balanced by a resilience to perturbations in network connectivity^36^ and an increased efficiency of cross-network communication.^5,37^ However, computational simulations predict that disruptions to brain hubs and their connections have far-reaching and adverse effects on overall network function.^38–40^ These predictions are borne out by observations from anatomical lesion studies^41–43^ and evidence of network hub alterations across a diverse range of psychiatric, neurodevelopmental and neurological brain disorders.^41,44–46^

Comparative connectomic studies have revealed that the evolutionary expansion of association cortex in humans is accompanied by a corresponding increase in areal connectivity.^47,48^ While the shape of the brain imparts significant constraints on wiring distance,^49^ computational models suggest that hub locations cannot be explained by cortical geometry or wiring costs alone.^50,51^ Instead, hub locations appear grounded in the spatial confluence of genetically programmed molecular gradients during early brain development with spatial differences in developmental timing reflected by commonalities in cytoarchitecture and regional gene expression in core network hubs.^18,20,52–54^ Recently, studies have begun to characterise the genetic architecture of white matter connectivity, revealing a highly polygenic background.^20,55,56^ Genetic influences on brain connectivity are concentrated on network hubs and their respective connections, supporting a role of genes in shaping the network core.^20^ Together, the vulnerability of network hubs in neuropsychiatric disorders and the strong genetic influence on hub connectivity suggests that the foundation of structural hubs is critical to later brain function.^17,57^

Using diffusion MRI, we and others have shown that the structural core of the human connectome is assembled early in development.^58,59^ Many fundamental network properties are in place in the human brain from as early as 30 weeks gestation.^58–61^ The availability of evidence from this developmental period is limited but reveals a right-tailed nodal degree distribution in the neonatal connectome, with modular organisation and high-degree hub nodes located along the cortical midline, insula and in lateral frontal and parietal cortex, forming a core enriched for long-range connections.^57–60,62^

In this study, we combine diffusion MRI acquired shortly after birth^63^ with a spatial atlas of the prenatal brain transcriptome^64–66^ to examine the organising principles of early structural brain networks. By isolating a molecular signature of network connectivity in the prenatal brain, we highlight the role of specific cell populations in the organisation and maintenance of early cortical circuitry and identify putative genetic risk factors for the disruption of developing network structure and brain growth.

## Results

### Highly-connected hub nodes in the neonatal cortex

We generated cortical structural connectivity networks using diffusion MRI and whole-brain probabilistic tractography in 208 neonates born at term (101 females; median gestational age at birth [range] = 40^+1^ weeks [37-42^+2^]; median post-menstrual age at scan [range] = 41 [37^+3^ – 44^+5^] weeks).^63^ Network nodes were defined according to the μBrain cortical atlas, a recently developed 3D neuroimaging-transcriptomic atlas of the developing human brain.^64^ For this study, each cortical region of the μBrain atlas (n=29) was subdivided into similar-sized subparcels of approximately 90 vertices each (n=349 per hemisphere, mean number of vertices ± S.D = 86.18 ± 14.02), yielding a high-resolution cortical parcellation with boundaries aligned to neonatal cortical anatomy (μBrain_90_; **Figure 1A**). Individual tractography streamlines were smoothed,^67,68^ combined to form a group consensus network and thresholded to include the strongest 15% of edges (**Figure 1B**).^20^ Alternative cortical subparcellations (μBrain_60;_ μBrain_120_) and network thresholds (5% and 25%) are explored in **Supplemental Materials** (**Figure S1-S2**).

**Figure 1:**
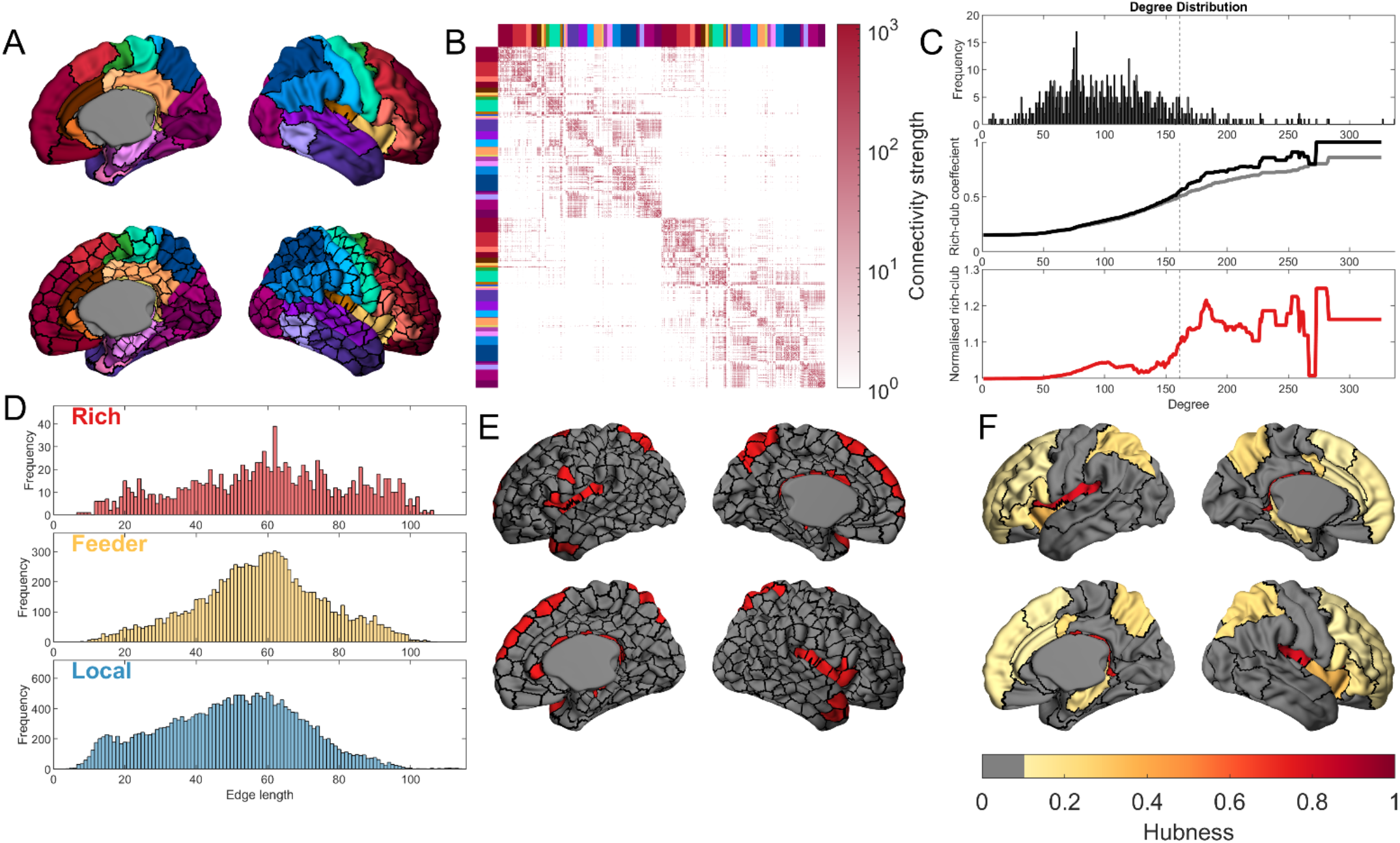
Rich-club hub nodes in the neonatal brain. **A)** The regions of the μBrain atlas (coloured regions; *top*) are subdivided into parcels of approximately 90 vertices (bottom). **B)** The group consensus connectivity matrix thresholded to retain the top15% of edges. Nodes are ordered according to the μBrain region of which they are a subset. **C)** The degree distribution (*top*) of the thresholded, group consensus network. The rich-club coefficient (*middle*) is calculated over all degree thresholds in the empirical data (black line) and compared to the rich-club coefficient of degree sequence preserving null networks (grey line). Normalised rich-club coefficient (*bottom*; red line) values > 1 indicate greater rich-club organisation than expected by chance. The dashed vertical line indicates the 90^th^ percentile for node degree (*k* = 161), above which nodes were considered network hubs. **D)** Edge length distributions for rich (connections between hub nodes), feeder (connections between hubs and non-hubs), and local (connections between non-hubs). **E)** The location of hub nodes (red) in the neonatal brain. **F)** The hubness for μBrain regions, calculated as the proportion of a region’s subparcels that were identified as a hub (hubness thresholded at 0.1 for visualisation).

Examining the properties of the neonatal connectome, we found that node degree distribution showed a characteristic heavy-tail, indicating the presence of a small population of nodes with disproportionately high degree (**Figure 1C**)^2,29^. We calculated the network rich-club coefficient *ϕ*(*k*) over increasing degree thresholds, *k*, and observed significant rich-club organisation emerging amongst the top 10% of connected nodes (*k* > 161, **Figure 1C**). Over all pairs of connected nodes, the median connection length was 54.6mm, with a higher proportion of long-range connections present between hub nodes (**Figure 1D**).^10,30,58,69,70^ Rich-club nodes were distributed across cortical areas including the insula, cingulate, ventrolateral frontal and dorsal parietal cortex (**Table S1**), mirroring earlier observations in neonatal^58,59^ and adult data^20,29^ and confirming the early establishment of a structural network core in the human brain (**Figure 1E-F)**. Repeating this analysis with each participant’s structural connectivity data revealed a consistent hub organisation was present across individual neonatal brain networks (**Figure S3**).

### Differential gene expression in neonatal network hubs prior to the time of birth

Cortico-cortical circuitry is established during the second trimester.^71–75^ We next sought to test whether the locations of high-degree network hubs are predicated on differences in gene expression in the mid-gestation cortex. Using microarray data from four post-mortem prenatal brain specimens (16-21 weeks post-conception)^65^ aligned to the μBrain cortical atlas,^64^ we tested associations between regional node degree and gene expression across five transient tissue zones of the fetal brain.

We identified 653 significant associations (581 unique genes; **Table S2**) between prenatal gene expression and the average node degree of each μBrain cortical parcel (p_FDR_<0.05; **Figure 2**) with most associations confined to postmitotic tissue compartments (cortical plate, subplate, intermediate zone; **Figure 2A-B**). In total, higher connectivity at birth was associated with increased expression of 293 genes (*hub*+) and decreased expression of 289 genes (*hub*−). Similar results were observed across different μBrain parcellation resolutions (**Figure S4**).

**Figure 2:**
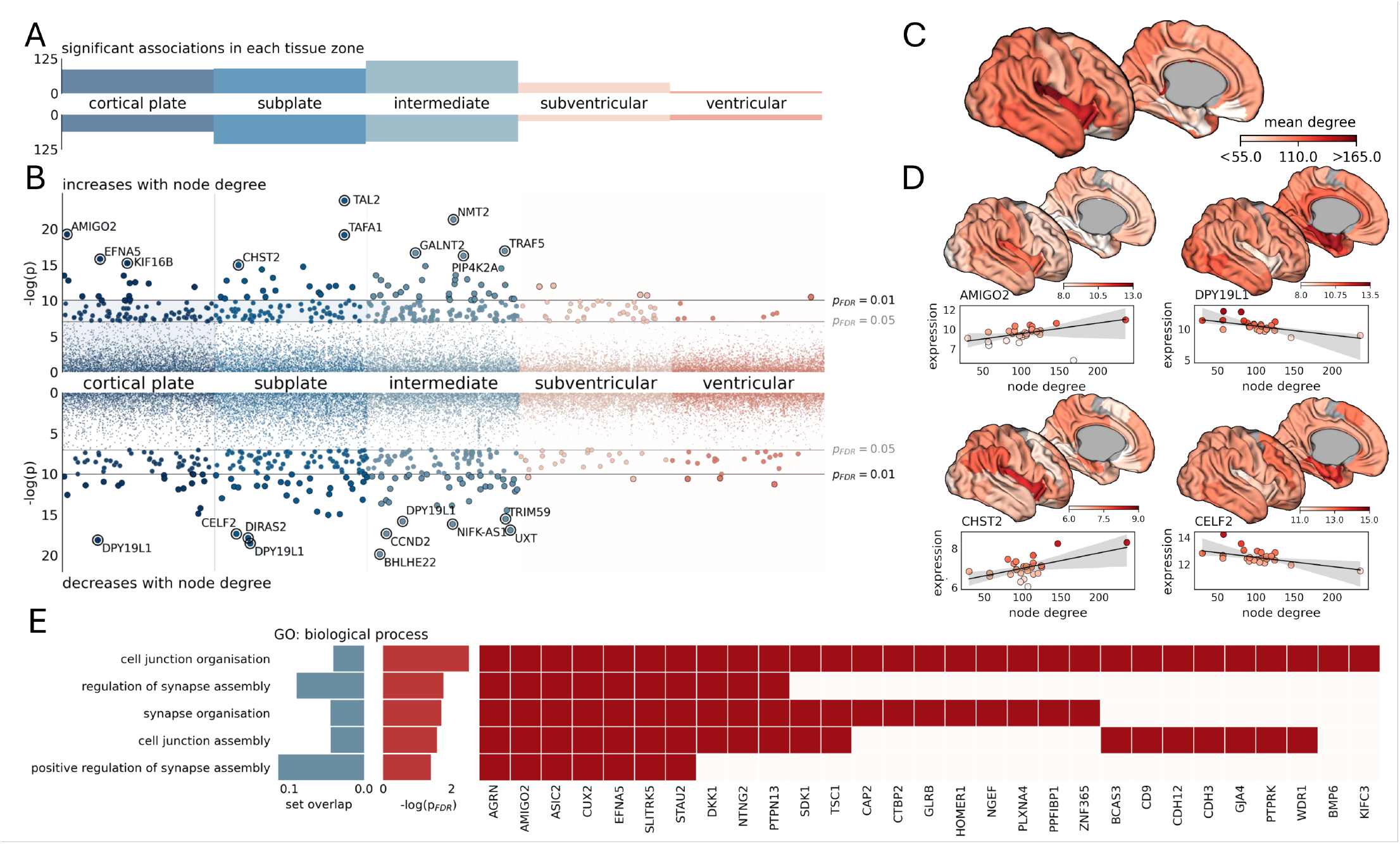
Genes expressed at mid-gestation in cortical regions with high network degree at birth are associated with the development of cortical circuitry. **A)** Number of significant associations (p_FDR_<0.05) between regional estimates of network degree at term-equivalent age and gene expression in mid-gestation in each of five developmental tissue zones. **B)** Individual positive (top) and negative (bottom) associations with node degree for each gene (n=7485 in each zone). The top 10 strongest associations are annotated in each row. **C)** Mean node degree of each region in the μBrain cortical atlas. **D)** Average (log-normalised) expression of four genes with positive (hub+, left) and negative (hub−, right) associations with node degree, displayed on the cortical surface. Scatterplots illustrate associations between degree and average gene expression, averaged over all brain specimens. **E)** Gene ontology (GO) enrichments (FDR-corrected) for biological processes in hub+ genes.

The hub+ geneset contained several transcription factors including: *CUX1* (subplate: β_degree_=0.0084, p=3×10^−6^) and *CUX2* (subventricular: β_degree_=0.0117, p=8×10^−4^), factors that regulate dendritic morphology of postmitotic neurons and proliferation of neuronal precursors in the SVZ;^76,77^ *NR4A2* (intermediate zone: β_degree_=0.0196, p=5×10^−6^), a specific marker of early-born subplate neurons,^78^ and *KLF6* (cortical plate: β_degree_=0.0059, p=6×10^−4^; subplate; β_degree_=0.0056, p=9×10^−5^; intermediate zone: β_degree_=0.0048, p=4×10^−4^), which enhances neurite outgrowth *in vitro* (**Table S2**).^79^ Other hub+ genes that were correlated with network degree included *AMIGO2*, encoding an adhesion molecule involved in dendritic arborisation^80^ (cortical plate: β_degree_=0.0124, p=4×10^−9^), *EFNA5*, encoding the axonal guidance molecule ephrin-A5^81^ (cortical plate: β_degree_=0.0119, p=1×10^−7^) and *CHST2*, critical for neuronal plasticity in the developing cortex^82^ (cortical plate: β_degree_=0.0067, p=1×10^−4^; subplate: β_degree_=0.0091, p=3×10^−7^; **Figure 2D**).

Hub− genes included transcription factors involved in neural progenitor fate determination including *EMX2* ^83^ (intermediate zone: β_degree_=-0.0104, p=3×10^−5^), *MYCL*^84^ (subplate: β_degree_=-0.0050, p=2×10^−6^), *SOX11*^85^ (subplate: β_degree_=-0.0036, p=7×10^−4^) and its downstream target *NEUROD1*^86,87^ (intermediate: β_degree_=-0.0150, p=9×10^−7^), as well as genes required for neuronal differentiation (*CELF2;*^88^ subplate: β_degree_=-0.0065, p=3×10^−8^) and radial migration of glutamateric neurons (*DPY19L1;*^89^ cortical plate: β_degree_=-0.0105, p=1×10^−8^; subplate: β_degree_=-0.0130, p=9×10^−9^; intermediate zone: β_degree_=-0.0031, p=5×10^−4^ ; **Figure 2D**).

Gene Ontology (GO) analysis with FUMA^90^ revealed significant enrichment of genes involved in synapse assembly and organisation in the hub+ geneset (**Figure 2E**) but no significant GO enrichments in hub− genes.

### Hub genes are expressed by excitatory neurons and astrocytic populations in mid-gestation

Using a comprehensive single-cell atlas of the pre- and postnatal human brain, we examined the expression of hub+ and hub− gene sets across different developmental cell lineages.^91^ We found that hub+ were enriched across excitatory neuron lineages in the cortical plate, subplate and intermediate zones, with specific enrichment in layer 5/6 intratelencephalic (L5/6-IT) and subplate (SP) neurons in the subplate and intermediate zones (p_FDR_<0.05**; Figure 3A-B**; **Table S3**). Hub+ genes in the cortical plate and subplate were also enriched in astrocyte cell lineages (p<0.05, uncorr). In contrast, hub− genes were not significantly enriched in any cell lineage after correction for multiple comparisons (**Table S3**).

**Figure 3:**
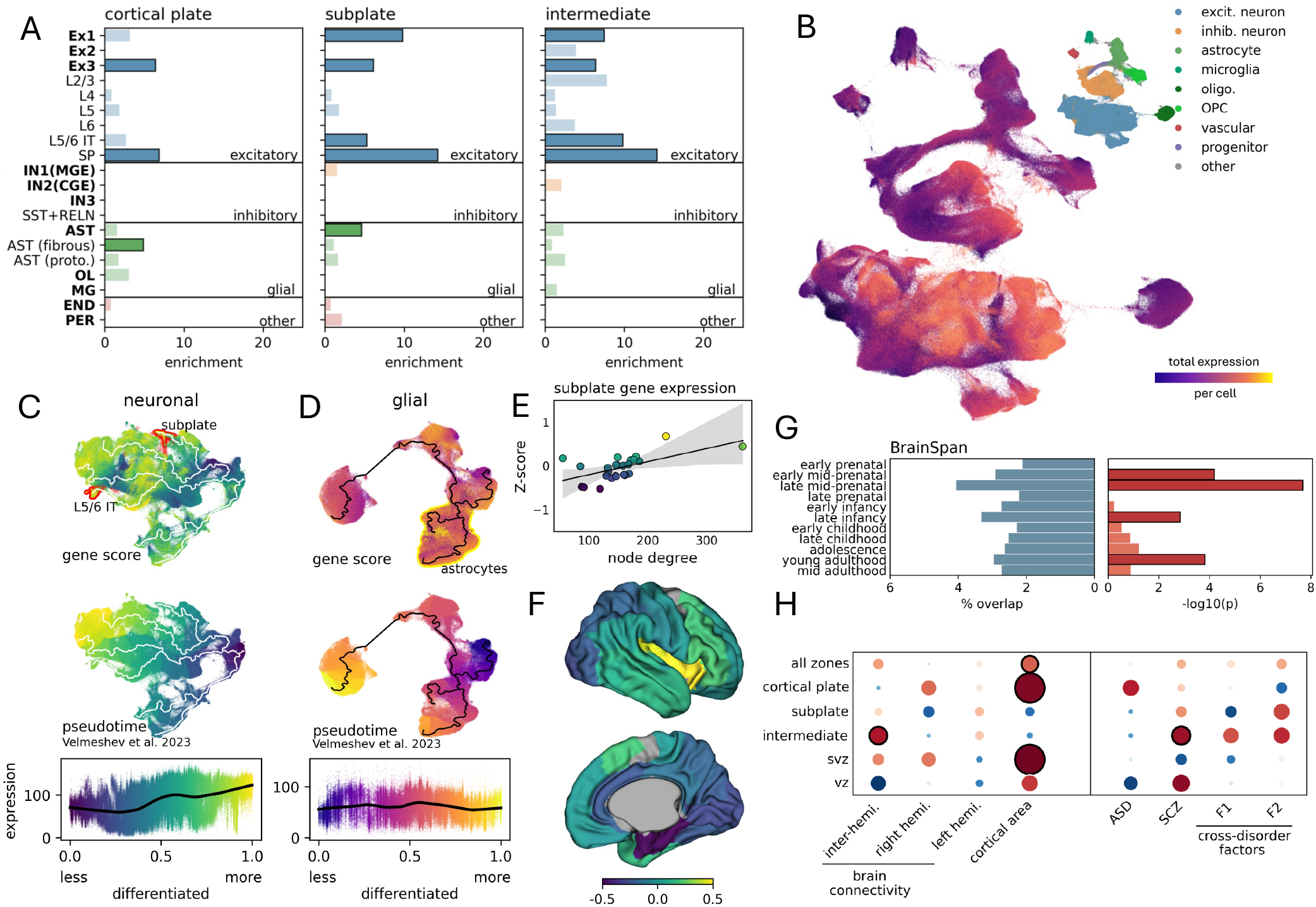
Genes with increased expression in highly connected nodes are enriched for subplate neurons and astrocytes. **A)** Enrichment of hub+ genes in developmental cell lineages (bold) and subtypes in each post-mitotic tissue zone. Significant enrichments (p<0.05 uncorr.) are highlighted with black outlines. **B)** Total (log-normalised) expression of network hub+ genes in each of 709,372 cells derived from brain tissue sampled across the human lifespan displayed using UMAP. **Inset**: Cell territories for major classifications. **C)** UMAP projections of neuronal and **D)** glial cell populations (top), overlaid with developmental pseudotime trajectories. Differentiated subplate neurons, layer 5/6 IT neurons and astrocyte territories are highlighted with red and yellow, respectively (top row). Plots are coloured by total gene expression of hub+ genes (top) and developmental time (middle). Scatterplots show relationship between hub+ gene expression and cell maturation for each population. **E)** Association between node degree and average expression (Z-scored) of hub+_SP_ genes. **F)** Average hub+_SP_ expression displayed on the μBrain cortical atlas (lateral and medial surfaces). **G)** Developmental enrichment of hub+ genes in the BrainSpan RNA-seq dataset. **H)** Enrichment in hub+ genes of SNPs associated with brain phenotypes^56,95^, neurodevelopmental disorders of early (ASD)^96^ and later (SCZ)^97^ onset and cross-disorder behavioural factors (F1: mood disorders and F2: psychopathology)^98^ (black circles indicate p<0.05 uncorr.).

Focusing on post-mitotic neuronal and glial populations, we examined the maturational state of cells expressing hub+ genes (**Figure 3C-D**). Using pseudotime estimates from single-cell expression profiles,^91^ we found that total hub+ gene expression increased with maturation in neuronal populations, with highest expression in differentiated subplate and layer 5/6 neurons. We confirmed that the regional expression of hub+ genes by subplate neurons (hub+_SP_) was spatially correlated to node degree (**Figure 3E-F**) and, through examination of independent bulk-tissue RNA-seq data from BrainSpan,^90,92^ that hub+ genes were enriched during the peak period of subplate development in mid-to-late gestation (**Figure 3G**).^93,94^ In glial cell populations, hub+ expression was less well-defined by maturation, with higher levels of expression in both glial progenitors and OPCs, as well as astrocytic cell populations (**Figure 3D**).

### Prenatal hub genes are enriched for human cortical expansion and areal connectivity strength in adulthood

Observations from comparative connectomic studies have revealed an evolutionary association between cortical expansion and areal connectivity across mammalian species.^47,48,99^ To test for potential shared mechanisms of human cortical wiring and expansion *in utero*, we compared hub+ and hub− gene sets with genes previously linked to the rate of human fetal cortical expansion,^64^ revealing significant overlap with hub+ genes expressed in the cortical plate (enrichment=4.29, p=0.007), subplate (enrichment=10.56, p<0.001) and intermediate zones (enrichment=12.87, p<0.001) and with hub− genes in the cortical plate (enrichment=5.96, p=0.001), subplate (enrichment=5.72, p<0.001), intermediate (enrichment=3.75, p=0.012) and subventricular zones (enrichment=6.01, p = 0.044; **Table S4**).

Overlapping genes included *CDH12* and *EMID1*, both with roles in cell adhesion and extracellular matrix organisation,^100,101^ *TRAF5* and *GNAO1*, involved in cell signaling^102,103^ and *CUX1*. We also examined associations with corresponding adult brain phenotypes using data from recent GWAS studies,^56,95^ identifying significant enrichment of loci associated with interhemispheric connectivity strength in hub+ genes in the intermediate zone (MAGMA: β = 0.18, p = 0.023) and with cortical surface area in hub+ and hub− genes in the cortical plate (β = 0.49, p = 0.001; β = 0.41, p = 0.012, respectively; **Figure 3H**; **Table S5**).

Compared to other primate species, genes differentially expressed in enlarged human cortex (hDEGs) are also enriched in pathways related to synaptic connectivity and are located near to genomic regions with higher substitution rates (human accelerated regions; HARs), deletions (human conserved deletions; hCONDELs) or rapid divergence from primate ancestors (human ancestor quickly evolved regions; HAQERs), suggesting adaptive changes that support increasingly complex brain networks between cortical areas in higher-order primates.^47,99,104–107^ We tested if prenatal hub genes were enriched for hDEGs located near to human-specialised genomic regions. We found that hub+ genes were enriched for genes located near to both HARs and HAQERs (all zones: enrichment = 1.86, p = 0.0037; enrichment = 2.93; p = 0.0004, respectively; **Table S4**). Tissue specific enrichments in hub+ gene sets were restricted to the cortical plate (HARs: enrichment = 2.60, p = 0.0055), subplate (HAQERs: enrichment = 4.96, p = 0.0003) and intermediate zones (HARs: enrichment = 2.28, p = 0.0073). Hub− genes were additionally enriched for HARs and HAQARS in the cortical plate (HARs: enrichment = 3.24, p = 0.018; HAQARS: enrichment = 3.51, p = 0.029).

Hub node connectivity is altered across a range of neurodevelopmental disorders.^41,45,46,108^ To assess the clinical relevance of hub gene expression in the prenatal brain, we examined the overlap between hub+ and hub− gene sets and single gene mutations linked to ASD (high confidence *SFARI* genes)^109^ and other neurodevelopmental disorders (NDD) characterised by brain malformations and/or cognitive sequelae (*Gene2Phenotype* category: definitive).^110^ While several high-confidence NDD genes were identified within hub gene sets, including *AFF2* (Fragile X-E syndrome, hub+), *GNAO1* (epileptic encephalopathy, hub+), *SMARCA2* (Nicolaides-Baraitser syndrome, hub+) and *RTTN* (polymicrogyria, hub−), we found no significant enrichment of pathogenic ASD or NDD variants in either hub+ or hub− genes outside of the ventricular zone, where 2 out of 8 hub+ genes (*TSC1, ANP32A*; enrichment = 15.78, p = 0.006) were identified (**Table S4**). We performed additional enrichment analyses using MAGMA^111^ across an array of previously published genome-wide association studies (GWAS; **Figure 3H**). We observed a weak enrichment of SNPs associated with ASD^96^ in prenatal hub+ genes expressed in the cortical plate (β=0.169, p=0.057; **Figure 3H**; **Table S5**) whereas both hub+ and hub− in the intermediate zone and cortical plate were enriched for schizophrenia^97^ (β=0.185, p=0.045; β=0.264, p=0.040, respectively). Additional weak associations were observed for hub+ genes in the intermediate zone for transdiagnostic symptoms of mood disturbance (F1; **Figure 3H**; β=0.153, p=0.057) and psychopathology (F2; β=0.166, p=0.055)^98^.

## Discussion

Cortico-cortical networks are critical for brain function.^1–5^ The foundations of macroscale brain networks: richly-connected hubs that support stereotyped patterns of connectivity between distributed cortical areas, are in place by the time of normal birth.^29,58,59^ In this study, we uncover a molecular signature of core network hubs captured in mid-gestation and enriched for developmental processes critical for the establishment and organisation of interareal circuitry.

Hub locations in the human brain network are conserved across development^112^ and consistent with those borne out by experimental tracer studies in non-human primates and other mammalian species. ^28,32,113^ During brain development, the differentiation of cortical areas is guided by morphogenic transcription factors along a spatiotemporal schema established early in gestation.^114–116^ Subsequently, the areal patterning of cytoarchitecture, axonal connectivity and function is reflected by the diversity of gene transcription across the cortex.^20,114,117,118^ Regional connectivity is both highly heritable and tightly coupled to cytoarchitectural and transcriptomic similarity in the adult brain,^20,56,117^ suggesting the locations of hub regions in cortical networks are constrained by genetic factors. We find that at mid-gestation, putative hub regions were associated with increased expression of genes supporting neuronal growth, synaptic plasticity and circuit formation compared to non-hub regions, which were characterised by genes linked to earlier progenitor and migratory processes (e.g., neural progenitor fate determination, radial migration). This pattern signals an accelerated developmental timeline in cortical hubs in place by mid-gestation, potentially allowing for an extended period of circuit formation.

Neurogenesis does not terminate across the cortex simultaneously, ending first in (para)limbic and allocortical structures before following a broadly rostral-caudal axis across the neocortex.^119–123^ Computational modelling reveals that this form of areal heterochrony can produce networks with similar properties to empirical brain networks, including high degree hub nodes.^54^ Thus, early completion of neurogenesis may afford hub nodes a prolonged period of circuit integration, following a ‘rich-get-richer’ wiring principle.^124^ Indeed, many hub regions display decreased neuronal density,^53,125^ a weak eulaminate structure,^126^ and denser dendritic arborization,^127^ cytoarchitectural properties that imply the earlier completion of neurogenesis.^120,121,125,126^ As an example, the insula, identified here as a key network hub in the neonatal brain, is one of the first cortical areas on the lateral surface to mature.^128^ Though the insula lacks its own proliferative zone, neuron migration from the pallial-subpallial border occurs early in gestation,^129^ and by 20 weeks, synaptic density is higher in the insula than other cortical areas.^130^ Consequently, delta brushes – electrophysiological hallmarks of the maturing cortex – arise first from the insula, at around 30 weeks gestation.^131^

The early formation of hub circuitry may confer an additional advantage due to the high wiring cost associated with longer cortical connections. In line with earlier reports, we find that connections between hubs nodes are, on average, longer than those between non-hubs.^5,58^ During brain development, exuberant interareal outgrowth of axonal connections is followed by a period of refinement, with extraneous connections pruned to define the mature connectome.^132–134^ A head start forming connections, taking advantage of the compact size of the human brain earlier in gestation, would reduce the high metabolic costs of establishing critical long range connections through non-specific exuberant growth.^17,57,70,124^ The enrichment of hub genes in human-specialised regions of the genome further supports evidence of adaptative mechanisms in place that may mitigate the cost of long-range network connectivity in the expanded human brain.^18,132^

Due to the inherent challenges of acquiring high-quality fetal diffusion MRI data,^133–136^ few studies have examined the emergence of structural brain network properties *in utero*,^137^ with most studies relying on the *ex utero* examination of preterm infants.^58–60^ In a recent example, Chen et al. used diffusion MRI to define cortico-cortical networks from 26 weeks, revealing a significant increase in connectivity over the second and third trimester in putative hubs including the cingulate and superior parietal cortex.^137^ However, due to acknowledged difficulties in fetal acquisition and processing, interhemispheric connections were excluded from the analysis yielding an, as yet, incomplete picture.^137^ Focusing on individual tracts, postmortem anatomical studies have demonstrated that the path of major commissural and projection fibers can be traced from mid-gestation with diffusion MRI,^74,75,138^ with similar results recently reported *in vivo*.^135^ But, while the organisation of immature axons into major white matter bundles is evident from as early as 10 weeks, synapses do not form in the developing cortical plate until after 20 weeks^139^ with axon terminals accumulating first in the transient subplate ‘waiting zone’.^71,93,94^

Subplate neurons are among the earliest born and maturing cells of the cerebral cortex,^78^ settling subjacent to the cortical plate to facilitate the emergence of neuronal circuitry and assist with the guidance of intratelencephalic and thalamocortical axons to their final cortical targets.^71,93,94,140^ Here, we find that genes expressed early in densely connected hub nodes are enriched in differentiated subplate neurons, including the canonical marker *NR4A2*, ^78^ as well as early differentiating, intratelencephic deep layer (L5/6) neurons. Similarly, we identified a moderate enrichment of hub+ genes in astrocytes, key participants in early neural circuit formation.^141^ Subplate neurons are critical to the establishment of thalamocortical connections,^93^ with their removal preventing the functional maturation of the cerebral cortex.^142^ As well as their prominent role guiding thalamic innervation, subplate neurons also connect to each other over long-distances, even across hemispheres,^143–146^ thus providing an initial substrate for cortico-cortical communication. Our findings suggest that, from as early as 15 weeks gestation, putative cortical hubs can be characterised by the transcriptional signature of circuit formation and synaptic assembly in the developmental subplate. In humans, the subplate in association areas is thicker and persists longer into gestation compared to other regions,^142,143^ this elongated period of developmental circuit formation may be vital to the increased complexity and capacity of hub areas in human brain networks.^144^

We identified several hub genes with pathogenic variants linked to neurodevelopmental disorders, including *AFF2, GNAO1* and *RTTN* but only weak enrichment of GWAS loci associated with ASD and SCZ. Hub dysconnectivity has been identified as a potential substrate for a number of brain disorders, suggesting a potential developmental vulnerability that may result in abnormal neuronal circuitry. ^41,44–46^ Although these conditions are thought to partially emerge from atypical developmental processes, the lack of strong genetic associations in our findings may reflect that our data is only a snapshot of the development of network connectivity in the fetal brain, focused on the establishment of neural circuitry in mid-gestation. Indeed, other factors are clearly involved in shaping early network connections. One of the most notable is the innervation of thalamocortical connections. Thalamocortical connections have a key role in shaping the arealisation, connectivity, and cytoarchitecture of the cortex.^94,114,115,121,147^ To gain a comprehensive view of how genetics shape brain network organisation, the contributions of non-cortical regions should be considered by future work. Similarly, while our findings align with other evidence that hub areas follow a distinct developmental trajectory that begins earlier than in non-hub regions,^17,57,124^ gene expression varies markedly during development^92,148^ and a more comprehensive evaluation of temporal development of hub gene expression during this critical window is warranted when such data is available.

A further limitation of our work is that we based our assessment of hub connectivity on binary network topology. While the number of axonal connections between regions spans orders of magnitude,^24,149^ appropriately defining weights such that they reflect the underlying strength of axonal connectivity using diffusion weighted imaging metrics remains challenging.^150,151^ However, in brain networks, weighted and unweighted measures of hubness tend to coincide,^152^ we are therefore confident that binary topology is sufficient to identify the key network hubs in the developing brain.

Taken together, our findings reveal that the establishment of hub connectivity, vital for long-range integration across brain networks, is underway by mid-gestation and characterised by distinct, spatially patterned programs of gene expression *in utero*.

## Methods and Materials

### Participants

Participant data was acquired from the third release of the Developing Human Connectome Project (dHCP).^63^ Ethics approval was granted by the United Kingdom Health Research Ethics Authority, reference no. 14/LO/1169. The full cohort comprised 783 neonates (360 female; median birth age [range] = 39^+2^ weeks [23-43^+4^]) across 889 scans (median scan age [range] = 40^+6^ [26^+5^-45^+1^] weeks; 107 neonates were scanned multiple times). For this study, only neonatal scans acquired from term-born infants with a radiological score of 1 or 2 (indicating no/minimal radiological abnormalities or pathologies) that also met additional quality control for in-scanner motion criteria (see below) where included, resulting in a final cohort of 208 neonates (101 females; median gestational age at birth [range] = 40^+1^ weeks [37-42^+2^]; median post-menstrual age at scan [range] = 41 [37^+3^ – 44^+5^] weeks).

### MRI acquisition and processing

Images were acquired on a Phillips Achieva 3T scanner at St Thomas Hospital, London, United Kingdom using a dedicated neonatal imaging system.^63,153^ T2-weighted Fast Spin Echo (FSE) multislice images were acquired in sagittal and axial orientations with overlapping slices (TR = 12000 ms; TE = 156 ms; resolution = 0.8 × 0.8 × 1.6 mm, 0.8 mm overlap). Sagittal and axial image stacks were motion corrected and reconstructed into a single 3D volume.^154^ Diffusion MRI was acquired with a spherically optimised set of directions over 4 b-shells (20 volumes × b = 0 s/mm^2^; 64 directions × b = 400; 88 × b = 1,000; 128 × b = 2,600)^155,156^ with a multiband factor acceleration of 4, TR = 3,800 ms; TE = 90 ms; SENSE: 1.2 and acquired resolution of 1.5 mm × 1.5 mm with 3-mm slices (1.5-mm overlap) reconstructed using an extended SENSE technique into 1.5 × 1.5 × 1.5 mm volumes.^157,158^

Structural images were processed using the dHCP’s minimal preprocessing pipeline, including bias correction, brain extraction, tissue segmentation and cortical surface reconstruction.^159^ Diffusion data were processed using a neonatal-specific pipeline.^160^ This included correction of susceptibility and eddy-current-induced distortions, motion artefacts and signal dropout,^161–164^ automated extraction of quality control (QC) metrics and nonlinear alignment to the 40-week dHCP neonatal template via each subject’s corresponding anatomical data^160,165–168^

### In-scanner motion quality control

To ensure only high-quality scans without major motion-related artefacts were used, we examined the quality control summaries of the dHCP diffusion processing pipeline.^160^ Scans more than two standard deviations away from the mean on any of the volume-to-volume motion, within-volume motion, susceptibility-induced distortions, and eddy current-induced distortions metrics were excluded from further analysis.

### Connectome reconstruction

Following pre-processing and QC, each subject’s diffusion data was used to generate whole-brain tractography following a previously developed processing pipeline in MRtrix3 (v3.0.2).^68,169^ Using 0 and 1000s/mm^2^ b-value diffusion volumes, we estimated a white matter response function from the oldest 20 term neonatal scans using the *dhollander* algorithm.^170,171^ Using this neonatal response function, fibre orientation distributions were calculated for each participant using single-shell 3-tissue constrained spherical deconvolution (CSD).^171^ Probabilistic whole-brain tractography was performed using second-order integration over fibre orientation distributions (iFOD2)^169^ (0.75mm step size; 45° maximum angle; 0.05 fibre orientation distribution cut-off), with Anatomically Constrained Tractography ^172^ to ensure the tracts followed biologically plausible pathways.

The parcels of the μBrain parcellation vary substantially in size, thus to limit potential bias due to large variations in surface area between nodes,^173^ we divided each region of the μBrain parcellation into smaller subparcels with an approximately equal number of vertices (n=90), yielding the μBrain_90_ parcellation with a total of n=698 cortical subparcels. To test the impact of network resolution, we created additional divisions of subparcels with an average of 60 (μBrain^60^; 1032 subparcels) and 120 (μBrain^120^; 528 subparcels) vertices.

To generate brain networks from the whole brain tractograms, we first applied Connectome Spatial Smoothing (CSS).^67,68^ CSS smooths streamline counts across vertices of a cortical mesh to improve reliability and robustness of individual whole-brain tractograms.^67^ To create a cortical mesh for CSS, the dHCP neonatal 40-week white matter surface was aligned to each individual’s diffusion space using transforms provided by the dHCP. Streamlines were assigned to the nearest cortical vertex of this mesh within a 5mm radius of their endpoint, with a Gaussian smoothing kernel applied (3mm FWHM, 0.01 epsilon) to adjust the strength of connectivity across cortical vertices, creating a high-resolution connectome. The high-resolution connectome was mapped to the μBrain_90_ parcellation by summing smoothed streamline counts over all vertices within each subparcel.

### Hub definition

Hubs were defined as previously described in adult data.^20^ We initially constructed a group consensus network by selecting connections that were (i) present in over 30% of individual networks, and (ii) in the top 15% of connections by strength.

To identify hubs within the consensus network, the rich-club coefficient was calculated across *k* degree thresholds as:

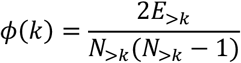

where *N*_>*k*_ is the number of nodes with degree > *k*, and *E*_>*k*_ is the number of edges between nodes with degree > *k*. As nodes with a higher degree are more likely to be connected to each other by chance, we generated 1000 random networks computed the rich-club coefficient *ϕ*_*rand*_(*k*) for each of these networks. The random networks were created by rewiring each edge of the group consensus network 50 times while retaining the degree sequence of the original network. The normalised rich-club coefficients were calculated as the ratio between the group consensus rich-club coefficient and the mean rich-club coefficient across the random networks for a given degree threshold, *k*:

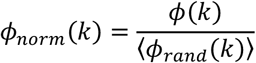

Values of *ϕ*_*norm*_(*k*) > 1 indicating that high-degree nodes are more densely interconnected than would be expected by chance, revealing the presence of rich-club organisation. Statistical significance is evaluated by calculating a p-value using the empirical null distribution of *ϕ*_*rand*_(*k*), derived from 1000 randomized networks.

Inspection of the rich-club coefficient curves indicated that high degree nodes form a rich-club regime beginning at *k* > 161. Therefore, we designated all nodes with degree >161 as hubs. To map hubs to the original μBrain parcellation, we generated an areal ‘hubness’ index defined by the proportion of subparcels within a given μBrain region designated as hubs, termed *RC*_%_.

### Transcriptomic data processing

Prenatal microarray data was made available as part of the BrainSpan database [https://www.brainspan.org/]. For details on tissue processing and dissection see Miller et al.^65^ Normalised microarray data were obtained from 1206 tissue samples across the left hemisphere of four post-mortem fetal brain specimens (age 15-21 PCW, 3 female).^65^ As described in prior work, each tissue sample location was matched to corresponding regional labels and tissue zones (cortical plate, subplate, intermediate zone, subventricular zone, ventricular zone) as part of the μBrain atlas.^64^ Samples that were not matched to labeled regions, including samples from subcortical nuclei, midbrain structures, subpial granular and marginal layers and brainstem were removed. Low signal probes designated ‘absent’ were also removed (34.67% of probes). Where multiple probes mapped to a single gene, the probe with the highest differential stability (DS) was assigned.^174^ We calculated DS as the average pairwise correlation of sample expression within each tissue zone and between pairs of specimens sampled at the same timepoint. Probes with DS<0.3 were removed.

Where more than one sample was available for a given region or zone, e.g.: samples from the outer and inner cortical plate in the same region, gene expression was averaged across samples. Finally, any probes with missing data in more than 10% of tissue samples were removed (n=2271). This resulted in expression data from 7457 genes across 27 regions and 5 tissue zones for analysis.

### Single-nucleus RNA data

Harmonised and log-normalised single-nucleus RNA sequencing profiles for 709,372 cells were downloaded from the UCSC Cell Browser (cells.ucsc.edu/?ds=pre-postnatal-cortex).^91^ Cell were sampled from 106 post-mortem brain tissue samples aged from approximately 16 gestational weeks to 54 years. Additional data included cell lineage gene markers, UMAP projections and pseudotime trajectories. For full details, refer to Velmeshev et al.^91^

### Statistical analysis

For each gene, we used a General Linear Model (GLM) to test the hypothesis that mid-gestation gene expression was associated with node degree (total number of connections) in each cortical region. GLMs were performed for each gene in each of 5 tissue zones (total number of tests = 7457 per zone) including the age of each specimen (15/16 or 21PCW) as a covariate. Robust linear models were fit to expression data using iteratively reweighted least squares to account for potential outliers. Significant associations between expression and node degree were identified after multiple comparison correction for False Discovery Rate within each tissue zone (p_FDR_<0.05)

### Enrichment analyses

GO enrichment was performed using *gene2func* implemented in FUMA.^90^ SNP enrichment was performed using MAGMA^111^ and summary statistics from recent GWAS studies.^56,95–98^ For other enrichment analyses, we calculated enrichment as the ratio of the proportion of genes-of-interest and proportion of background genes within each geneset. Unless otherwise stated, the background set was defined as the full list of genes included in the study (n=7457).

Significance was determined using the hypergeometric statistic:

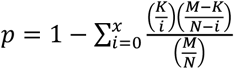

Where *p* is the probability of finding *x* or more genes from a specific geneset *K* in a set of randomly selected genes, *N* drawn from a background set, *M*.

## Acknowledgements

This research was supported by the Brain and Behavior Research Foundation (31471 to S.O.), an NHMRC Investigator Grant (1194497 to G.B.), Murdoch Children’s Research Institute, The Royal Children’s Hospital and the Victorian Government’s Operational Infrastructure Support Program. The project was generously supported by The Royal Children’s Hospital Foundation devoted to raising funds for research at The Royal Children’s Hospital.

## Author contributions

Conceptualisation, Methodology, Software, Analysis, Writing, Revision: S.O. & G.B. Supervision, Resources: G.B.

## Data and code availability

Neuroimaging data for the Developing Human Connectome Project are available via the NIMH Data Archive (collection ID: 3955). The μBrain atlas and associated data is available at: https://zenodo.org/records/10622337. Supporting code for this manuscript will be made available on github.com.

