## Supplemental Information for "Transcriptomic divergence of network hubs in the prenatal human brain"

S. Oldham[1] & G. Ball[1,2]

1. Developmental Imaging, Murdoch Children's Research Institute
2. Turner Institute for Brain and Mental Health, Monash University
3. Department of Paediatrics, University of Melbourne

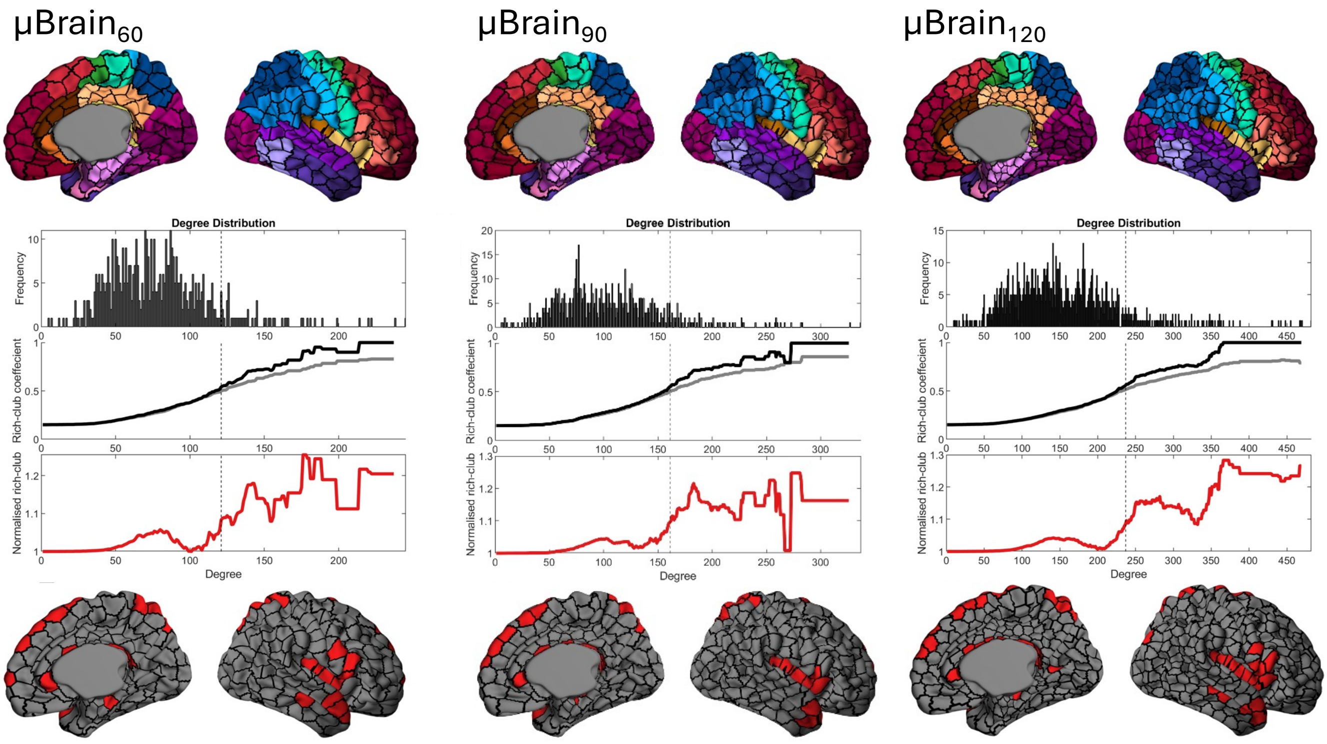


**Figure S1. Alternative cortical parcellations.** Cortical subdivisions (top row) are shown at three different resolutions (µBrain$_{60}$, µBrain$_{90}$ and µBrain$_{120}$) with subparcels of approximately 60, 90 and 120 vertices in area, respectively. Corresponding degree distributions and rich club coefficient curves (middle row) are shown for each parcellation resolution. The rich-club coefficient is calculated over all degree thresholds in the empirical data (black line) and compared to the rich-club coefficient calculated in degree sequence preserving null networks (grey line). Normalised rich-club coefficient (red line) values > 1 indicate greater rich-club organisation than expected by chance. The dashed vertical line indicates the 90$^{th}$ percentile for node degree in each parcellation to define network hubs. The location of hub nodes (bottom row, red) are shown overlaid on each parcellation resolution.

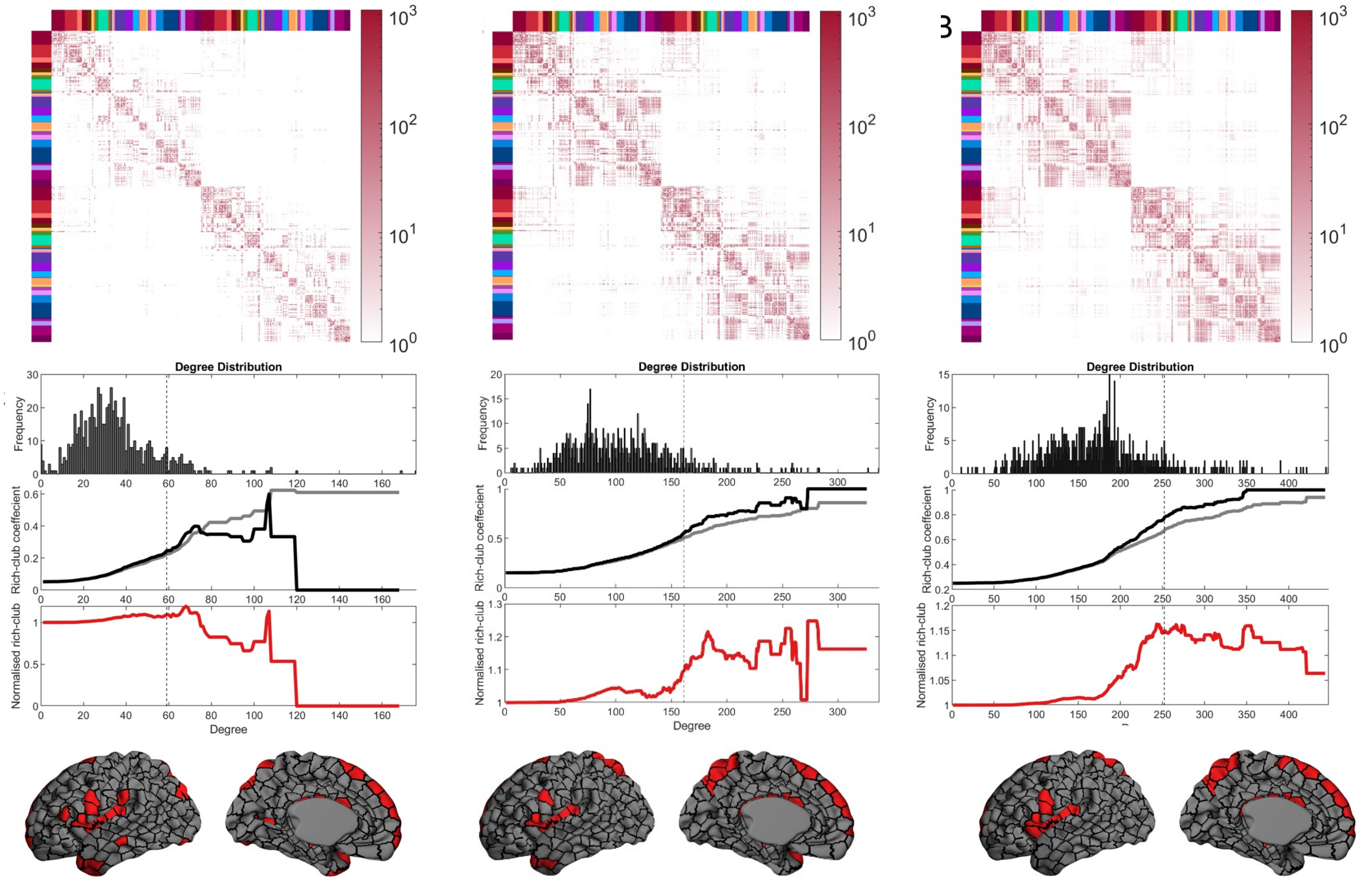


**Figure S2. Alternative network thresholds.** Group consensus connectivity matrices thresholded at three different levels (retaining the strongest 25%, 15% and 5% of all edges) are shown (top row) for the μBrain$_{90}$ parcellation. Corresponding degree distributions and rich club coefficient curves (middle row) are shown for each thresholded consensus network. The rich-club coefficient is calculated over degree thresholds in the empirical data (black line) and compared to the rich-club coefficient calculated in degree sequence preserving null networks (grey line). Normalised rich-club coefficient (red line) values > 1 indicate greater rich-club organisation than expected by chance. The dashed vertical line indicates the 90$^{th}$ percentile for node degree to define network hubs. The location of hub nodes (bottom row, red) in each consensus network are shown overlaid on the corresponding cortical parcellation.

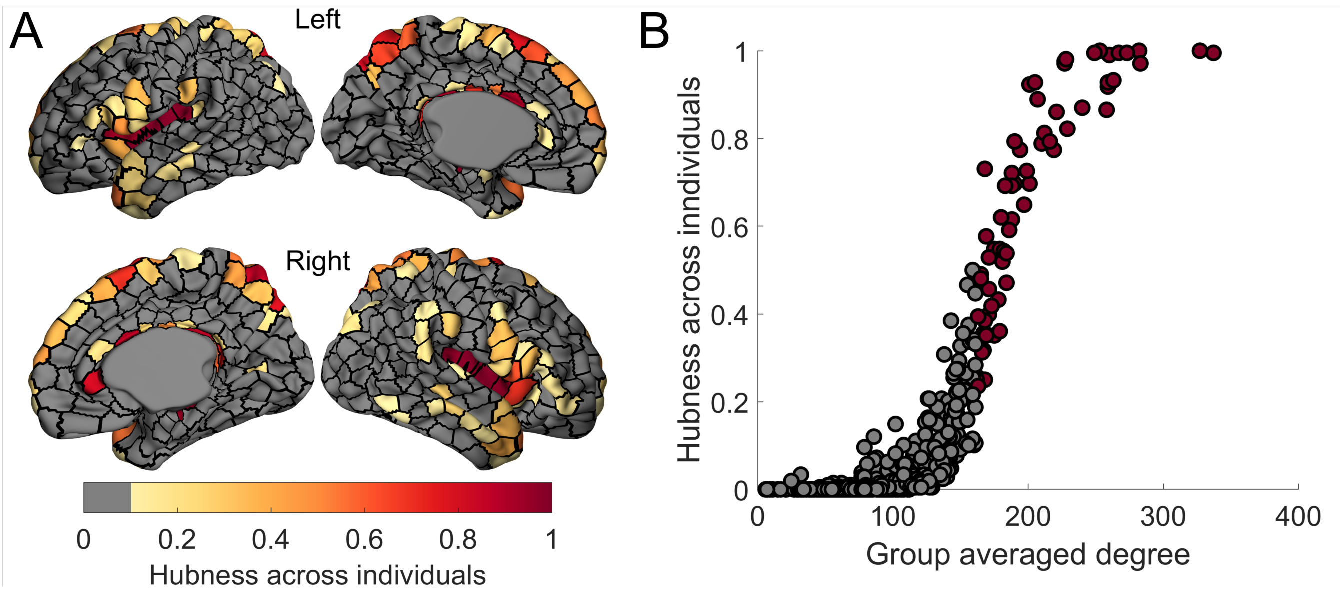


**Figure S3. A)** Hubness consistency across individuals. For each individual neonates μBrain$_{90}$ structural network (thresholded to the top 15% strongest connections), we identified hub nodes (90$^{th}$ percentile for degree). We then calculated the proportion of times a node was identified as a hub across individuals (hubness across individuals). **B)** Individual hubness plotted against the group averaged degree. Points in red were identified as a hub in the group averaged consensus network (90$^{th}$ percentile for degree).

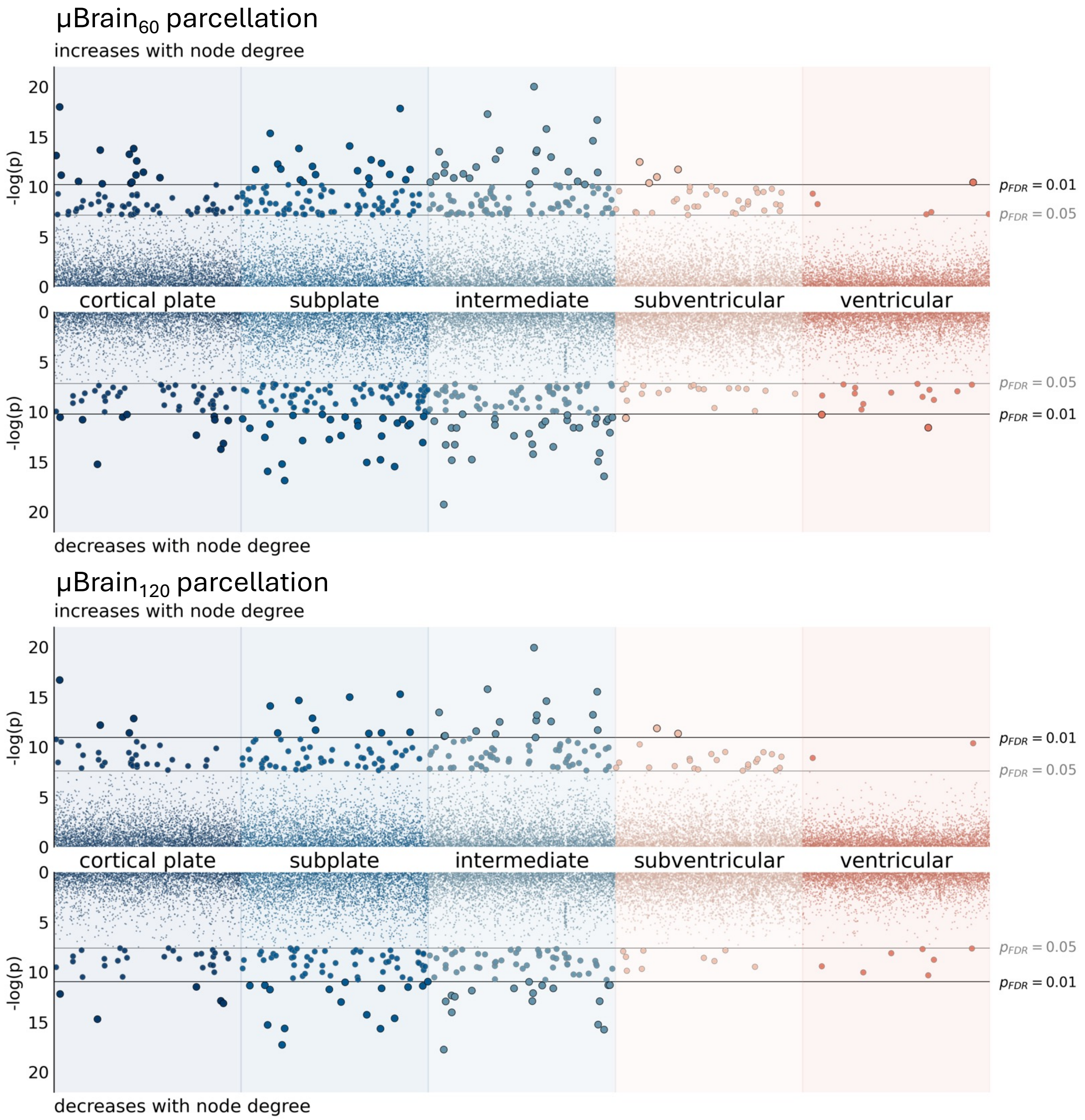


**Figure S4: Significant associations between node degree and gene expression at different network resolutions**
